# Inferring Disease-Associated Piwi-Interacting RNAs via Graph Attention Networks

**DOI:** 10.1101/2020.01.08.898155

**Authors:** Kai Zheng, Zhu-Hong You, Lei Wang, Leon Wong, Zhan-Heng Chen, Han-Jing Jiang

**Affiliations:** School of Computer Science and Technology, China University of Mining and Technology, Xuzhou, 221116, China; Xinjiang Technical Institutes of Physics and Chemistry, Chinese Academy of Science, Urumqi 830011, China

## Abstract

**Motivation:** PIWI proteins and Piwi-Interacting RNAs (piRNAs) are commonly detected in human cancers, especially in germline and somatic tissues, and correlates with poorer clinical outcomes, suggesting that they play a functional role in cancer. As the problem of combinatorial explosions between ncRNA and disease exposes out gradually, new bioinformatics methods for large-scale identification and prioritization of potential associations are therefore of interest. However, in the real world, the network of interactions between molecules is enormously intricate and noisy, which poses a problem for efficient graph mining. This study aims to make preliminary attempts on bionetwork based graph mining.

**Results:** In this study, we present a method based on graph attention network to identify potential and biologically significant piRNA-disease associations (PDAs), called GAPDA. The attention mechanism can calculate a hidden representation of an association in the network based on neighbor nodes and assign weights to the input to make decisions. In particular, we introduced the attention-based Graph Neural Networks to the field of bio-association prediction for the first time, and proposed an abstract network topology suitable for small samples. Specifically, we combined piRNA sequence information and disease semantic similarity with piRNA-disease association network to construct a new attribute network. In the experiment, GAPDA performed excellently in five-fold cross-validation with the AUC of 0.9038. Not only that, but it still has superior performance compared to methods based on collaborative filtering and attribute features. The experimental results show that GAPDA ensures the prospect of the graph neural network on such problems and can be an excellent supplement for future biomedical research.

**Supplementary information:** Supplementary data are available at Bioinformatics online.

## 1 INTRODUCTION

Piwi-interacting RNA (piRNA) is a small, non-coding RNA that clusters at transposon loci in the genome and is typically 24–32 nucleotides in length. Its discovery has greatly expanded the RNA world (Aravin, et al., 2007; Grimson, et al., 2008; Iwasaki, et al., 2015; Malone, et al., 2009; Yin and Lin, 2007). Since the discovery and formal definition of piRNA in 2006, the PIWI–piRNA field has been developed rapidly, and its functions in developmental regulation, transposon silencing, epigenetic regulation, and genomic rearrangement are being revealed gradually (Armisen, et al., 2009; Leslie, 2013; Marcon, et al., 2008; Pall, et al., 2007). piRNA interact exclusively with PIWI proteins which belong to germline-specific subclade of the Argonaute family (Moyano and Stefani, 2015). The best-known role of it is to repress transposons and maintain germline genome integrity through DNA methylation, as the depletion of PIWI leads to a sharp increase in transposon messenger RNA expression (Brennecke, et al., 2007; Siomi, et al., 2011). Compared with microRNA(miRNA) and small interfering RNA(siRNA) that are small RNAs, (1) longer than miRNA or siRNA; (2) only present in animals; (3) more diverse sequences and constitute the largest class of noncoding RNA; (4) testes-specific (Houwing, et al., 2007; Leslie, 2013; Moazed, 2009; Rajasethupathy, et al., 2012; Siomi, et al., 2011).

Recently, emerging evidence suggests that piRNA and PIWI proteins are abnormally expressed in various cancers (Assumpcao, et al., 2015; Cheng, et al., 2011; Chu, et al., 2015; Ng, et al., 2016; Romano, et al., 2017; Zou, et al., 2016). Therefore, the function and potential mechanism of piRNA in cancer become one of the important research directions in tumor diagnosis and treatment. For example, Fu et al. found that abnormal expression of piR-021285 promoted methylation of ARHGAP11A at the 5’-UTR/first exon CpG site, thereby promoting mRNA apoptosis and inhibiting apoptosis of Breast cancer cells (Fu, et al., 2015). Subsequently, Tan *et al*. found that down-regulation of piRNA-36712 in breast cancer increases SLUG levels, while P21 and E-cadherin levels were reduced, thereby promoting the malignant phenotype of cancer (Tan, et al., 2019). piR-30188 binds to OIP5-AS1 to inhibit glioma cell progression while low expression of OIP5-AS1 reduces CEBPA levels and promotes the malignant phenotype of glioma cells which discovered by liu *et al*. (Liu, et al., 2018). Also glioblastoma, Jacobs et al. found that piR-8041 can inhibit the expression of the tumor cell marker ALCAM / CD166, with the clinical role of targeted therapy(Jacobs, et al., 2018). In addition, piRNA is directly or indirectly involved in the formation of liver cancer. In 2016, Rizzo et al. found that hsa_piR_013306 accumulates only in hepatocellular carcinomas (Rizzo, et al., 2016).

piRNA is gaining enormous attention, and tens of thousands of them have been identified in mammals and are rapidly accumulating. In order to accelerate research in this field and provide access to piRNA data and annotations, multiple databases such as piRNA-Bank (Sai Lakshmi and Agrawal, 2007), piRBase (Wang, et al., 2018), piRNAQuest (Sarkar, et al., 2014) have been successively established. Subsequently, the role of piRNA and PIWI proteins in the epigenetics of cancer is constantly being discovered, and some of them can serve as novel biomarkers and therapeutic targets. Taking this as an opportunity, an experimentally supported piRNA-disease association database called piRDisease (Muhammad, et al., 2019) was proposed, which made it possible to predict potential associations on a large scale. Although many disease-related ncRNA prediction model have been proposed and gradually developed, predictors for disease-related piRNA is relatively unexplored (Li, et al., 2019; Wang, et al., 2019; Wang, et al., 2019; Zheng, et al., 2019; Zheng, et al., 2019).

In this paper, we propose a piRNA-disease association predictor based on attention-based graph neural network, called GAPDA. The study has three main contributions: (i) Introducing a graph neural network based on self-attention strategy, Graph Attention Network (GAT), which calculates the hidden representation of each node by attending over its neighbors. This GAT-based approach gathers the merits of representational learning and network-based approaches. (ii) An abstract network topology apply to small sample data is proposed. With the association as a node, it can expand the numerous heterogeneous network to replace the piRNA-disease association network. (iii) Different from traditional collaborative filtering and attribute-based methods, the proposed method integrates disease semantic information and piRNA sequence information, improves prediction accuracy and has higher coverage. On the association dataset piRDisease, GAPDA achieves an AUC of 91.45% with an accuracy of 84.49%. Compared with traditional methods, this method has higher precision. In general, the proposed method can provide new impetus for cancer mechanism research, provide new research ideas for small sample data sets, and determine the prospects of attention-based Graph Neural Networks on such issues. In addition, we hope that this work will stimulate more association prediction research based on graph neural network.

## 2 MATERIALS AND METHODS

### 2.1 Benchmark dataset

With the rapid increase of PIWI-interacting RNA (piRNA) related research, the contribution of piRNA in disease diagnosis and prognosis gradually emerges. These manually managed, complex and heterogeneous information may lead to data inconsistency and inefficiency, it put data analysis into a dilemma. To this end, the piRDisease database, which integrates experimentally supported association between piRNAs and disease, was proposed in 2019 (Muhammad, et al., 2019). Azhar et al. developed piRDisease v1.0 by searching more than 2,500 articles, which provided 7939 piRNA-disease associations with 4,796 piRNAs and 28 diseases. The baseline set by simple filtering is named GPRD, as shown in Table 1.

**Table 1.** The details of benchmark dataset GRPD

| Benchmark dataset | piRNA | Diseases | Associations |
| --- | --- | --- | --- |
| GPRD | 501 | 22 | 1212 |

#### GPRD

Currently, research on the relationship between piRNA and disease is in the ascendant, so the degree of some piRNAs are only 1 in the association network. Too many nodes with the degree=1 affect the performance of the network-based approach. Therefore, in GPRD, we only retained 501 piRNAs with the degree greater than 1 and constituted 1212 associations. The training dataset *T* can be defined as:

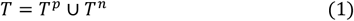

where *T^p^* is a positive subset of the piRNA-disease association construct in GPRD, and *T^n^* is a negative subset containing 1212 associations which were randomly extracted from all 11022 unconfirmed associations between piRNA and disease.

### 2.2 Construct new piRNA-disease association network

#### The structure of the network

At present, ncRNA-related associations with experimental verification are very limited, so the network-based method is difficult to achieve satisfactory prediction results. In addition, It’s difficult to get the desirable accuracy by attribute-based methods. In the meanwhile, biological data is often complex, and network representations computed from sparse networks cannot cover all real-world behavior information. Therefore, a method of enriching the hidden representations contained in sparse network is urgently needed. To this end, we propose a simple network construction method with association as a node. The new association adjacency matrix *A* based on *n* associations is calculated as follows:

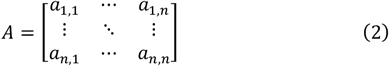

where *n* is the number of associations in the training dataset *T*. The element *a_i,j_* is set to 1 if the *i*-th association is related with the *j*-th association, otherwise 0. In particular, the links between associations is various. The process is shown in Figure 1. In this paper, we utlize piRNA and disease as link vectors, respectively, and define them as follows:

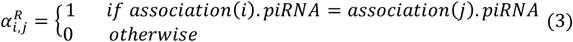

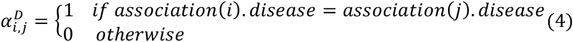

**Fig. 1.**
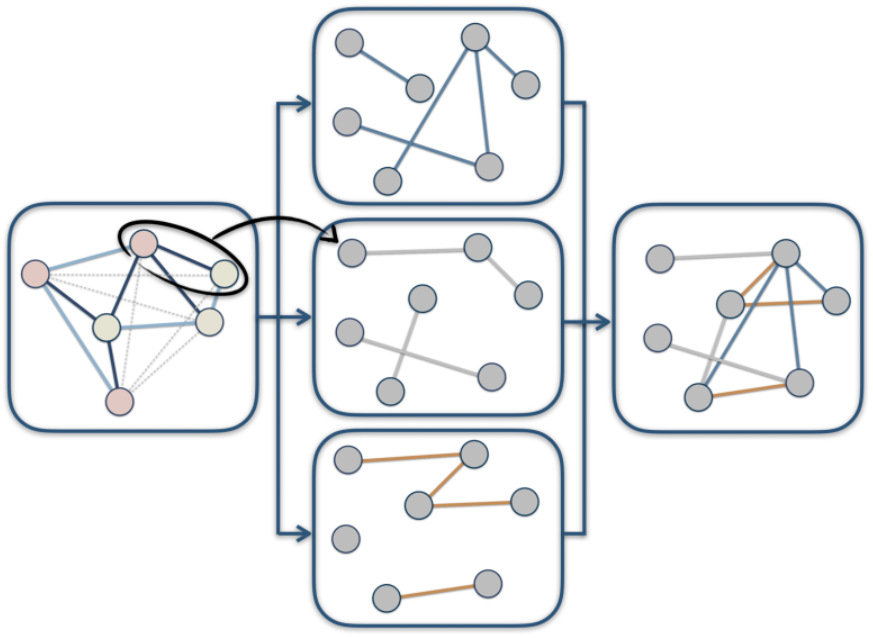
Explanation of the network reconstruction in node-level.

According to the above formula, a plurality of superimposable adjacency matrices can be obtained to enrich the structural information of the network, like *A^R^* composed of 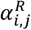 and *A^D^* composed of 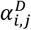. Since the size of the abstracted adjacency matrix is uniform, they can be stacked by weighting. For the sake of simplicity, we only performed a addition operation on the adjacency matrix *A^R^* and the adjacency matrix *A^D^*. Therefore, the element 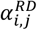 of the adjacency matrix *A^RD^* is calculated as follows:

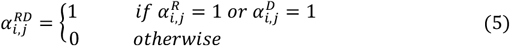

#### Node attributes

The attribute of the node is mainly composed of two parts: piRNA sequence features and disease semantic features. These two attribute information are described in detail below. The specific structure and function of RNA is determined by the sequence carrying the genetic information, so describing the sequence as a descriptor is an effective way to characterize its function. *k-* mers is a common alignment algorithm that the basic principle is to divide a sequence into sub-sequences of length *k* and count its frequency. Recent studies show that ncRNAs of related function often have related *k*-mer contents (Kirk, et al., 2018). For example, 3-mer of piRNA can be expressed as CCC, CCG, …, GGG. Herein, the *k*-mer deconstructs and reconstructs the piRNA functional features to obtain piRNA descriptor *Feature*(*p_a_*) where *p_a_* is piRNA with serial number *a*. The process is shown in Figure 2.

**Fig. 2.**
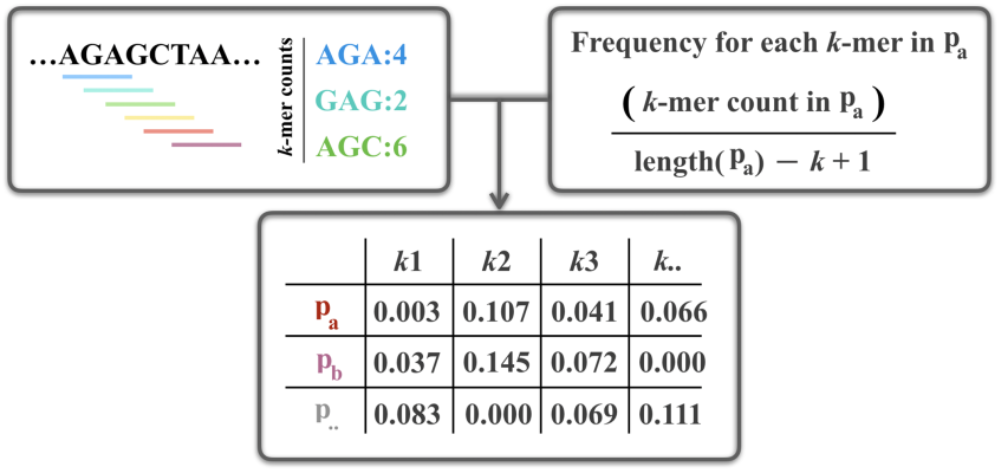
The flowchart for calculating piRNA sequence features.

It is still an urgent and tough problem to characterizate disease attributes. So far, methods for constructing directed acyclic graphs (DAG) by the Medical Subject Headings (MeSH) to quantify the relationship between diseases are commonly used (Xiang, et al., 2013). MeSH is the authoritative standard vocabulary produced by the National Medical Library. Because of its strict classification of diseases, it can deconstruct the semantic relationship of diseases. Taking Lip Neoplasms (LN) as an example (Figure 3), its MeSH ID is “C04.588.443.591.550; C07.465.409.640; C07.465.565.550”, and the corresponding parent nodes are Mouth Neoplasms and Lip Disease whose MeSH IDs are “C04.588.443.591; C07.465.565.550” and “C07.465.409.640”, as shown in Figure 3. Similarly, Mouth Neoplasms and Lip Disease also has their parent nodes, Mouth Disease and Head and Neck Neoplasms. According to the aforementioned analysis, Lip Neoplasms and other related diseases can be expressed as *DAG_LN_* = (*LN*, *T_LN_*, *E_LN_*), where *T_LN_* is a collection of nodes in *DAG_LN_* that contain LN, such as “Head and Neck Neoplams” and “Mouth Disease”. Furthermore, *E_LN_* is a collection of edges between different nodes, such as the edge between “Stomatognathic Disease” and “Mouth Disease.” Based on former research production (Xuan, et al., 2013), the semantic contribution *C* of disease *w* to disease *d* is calculated:

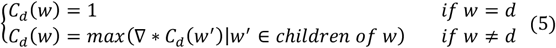

**Fig. 3.**
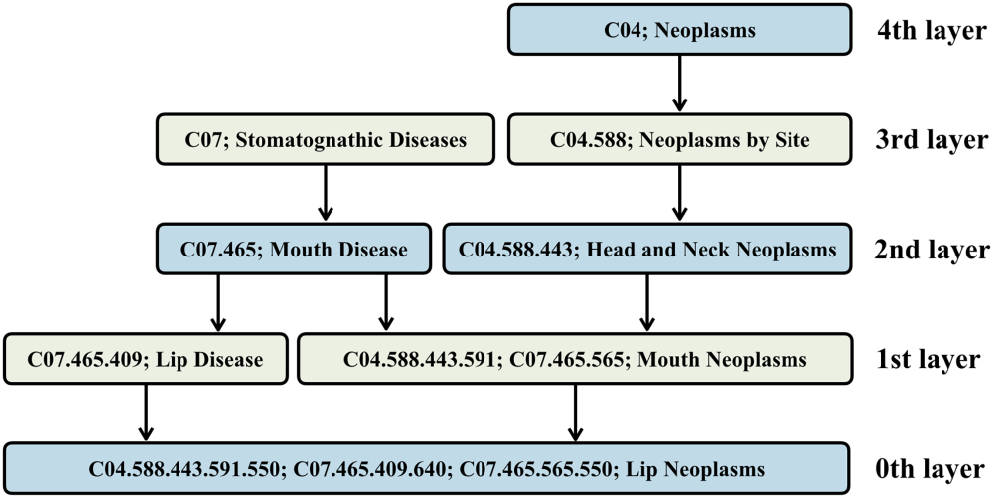
The directed acyclic graphs (DAG) of Lip Neoplasms.

**Fig. 4.**
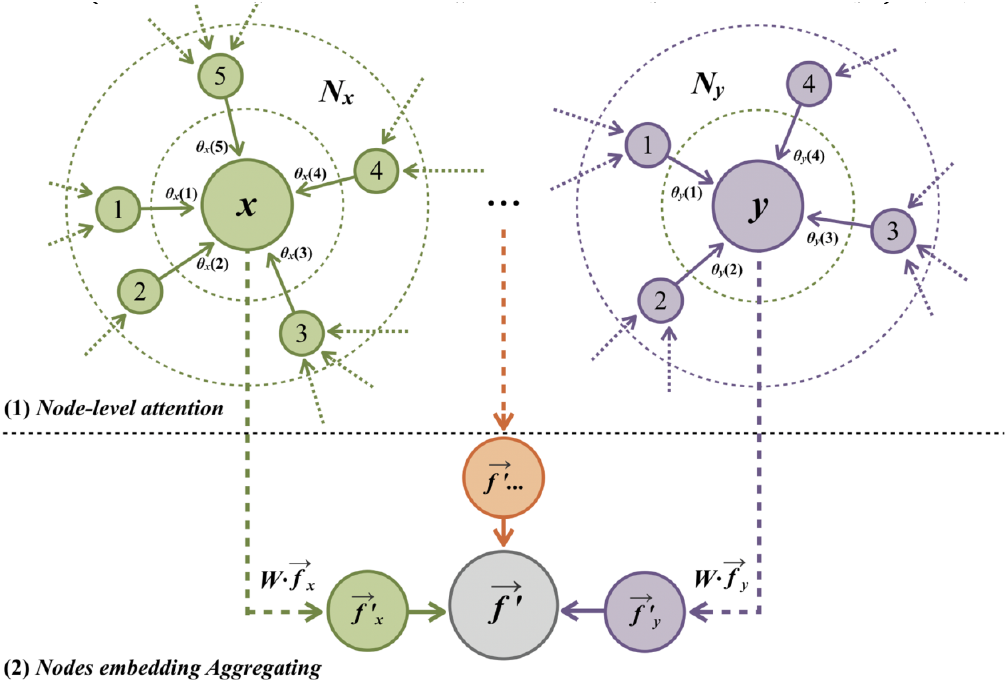
The flowchart for self-attention in node-level.

Here we set the seismic contribution decay factor ∇ to 0.5. *w*′ is the child node of *w*. If the disease *d* is farther apart from the disease *w* in the DAG, the contribution of the disease *w* to the disease *d* is lower. For example, “Neoplasms” contributes less to “Lip Neoplasms” than “Mouth Neoplasms”. According to the semantic contribution *C*, the semantic value *V* of disease *d* is calculated:

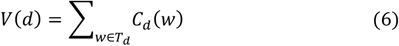

If the two diseases share more DAGs and near common ancestors, the two diseases are more semantically similar. Under that assumption, the semantic similarity scores *SS* for disease *a* and disease *b* can be defined as follows:

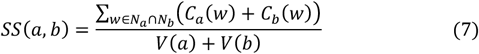

The semantic similarity score SS takes into account the existence of common ancestors between diseases. However, its performance is not unlimited. For example, “Neoplasms by site” appears in the DAGs of many diseases, while the “Stomatognathic Disease” of the same layer appears less frequently. Since “ Stomatognathic Disease “ has a higher specificity for “Lip Neoplasms”, its weight should also be higher. To quantify such differences in weight, the second semantic contribution is designed:

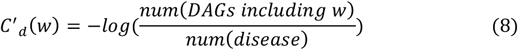

Similarly, the second semantic similarity scores *SS*′ for disease *a* and disease *b* can be defined as follows:

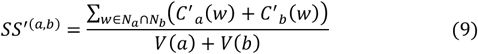

Both *SS* and *SS*′ are unilateral in principle. In order to combine the advantages of two semantic similarity scores, the comprehensive semantic similarity *S* is calculated:

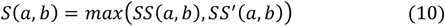

In this study, the degree of semantic association between disease *d_b_* and other diseases was used as the descriptor *Feature*(*d_b_*) for the disease.

### 2.3 Gaussian interaction profile kernel similarity

The Gaussian interaction profile kernel similarity(GIP) is a commonly used collaborative filtering algorithm. According to previous studies, the method can calculate the similarity matrix between ncRNAs and diseases from the known adjacency matrix (van Laarhoven, et al., 2011). In detail, the similarity between piRNA *p_a_* and piRNA *p_b_* can be defined as follows:

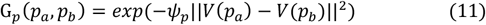

where *V*(*p_a_*) is a two-dimensional vector composed of the relationship between piRNA and all diseases, as is *V*(*p_b_*). In addition, *ψ_p_* as kernel width coefficient is defined as follows:

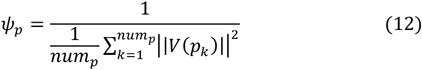

*num_p_* is the number of piRNAs. Similarly, the similarity between piRNA *d_a_* and piRNA *d_b_* can also be calculated by this algorithm:

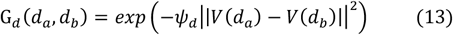

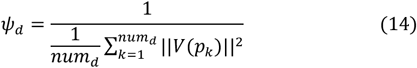

*num_d_* is the number of diseases. In this paper, it is compared as a traditional method with the proposed method. In this study, the degree of Gaussian association between piRNA *p_a_* and other piRNAs was used as the descriptor *Feature*′(*p_a_*) for the disease. And, the degree of Gaussian association between disease *d_b_* and other diseases was used as the descriptor *Feature*′(*d_b_*) for the disease.

### 2.4 Graph Attention Networks

Graph Attention Network (GAT) is a graph neural network based on self-attention mechanism proposed by Yoshua Bengio et al. in 2018 (Veličković, et al., 2017). The main contribution is to construct a hidden self-attention layer to specify different weights to different nodes in a neighborhood without any time-consuming matrix operations (such as inversion) or a priori knowledge of the graph structure. The input to the graph attention layer is *n* node features of length *H*, 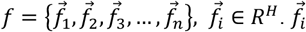 is the initial feature of the *i*-th node. And, the output of the layer is produced as 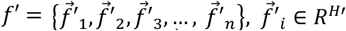, where *H* and *H*′ have different dimensions. 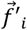 is the projected feature of the *i*-th node. In order to implement self-attention mechanism, a shared linear transformation parameter matrix *W* ∈ *R*^*H*′×*H*^ is designed to be applied to each node. Therefore, the attention coefficient *e_x_*(*y*) of node *x* to node *y* can be calculated as follows:

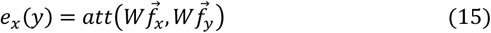

Here *att* denotes a mapping, *R*^*H*′^ × *R*^*H*′^ → *R*. It converts two vectors of length F′ into a scalar as the attention coefficient. In addition, self-attention assigns attention to all nodes in the graph, which obviously loses structural information. Therefore, a method called masked attention is proposed:

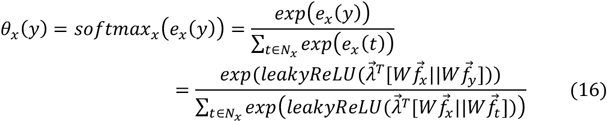

Where *N_x_* is the set of neighbor nodes of node *x*. *softmax_x_* is utlized to normalize the attention coefficient *e_x_*(*y*) to obtain the weight coefficient *θ_x_*(*y*). 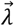 is the weight coefficient vector of the graph attentional layer, and the length is 2*F*′. *LeakyReLU* is the activation function. *T* represents transposition and ║ represents connection operation. Therefore, the embedding of node x can be fused by the projected node features of neighbors with different weights, as follows:

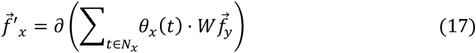

In order to solve the problem of large variance of the graph data caused by the scale-free of the heterogeneous graph, mult-head attention is performed to make the training process more stable. Specifically, features of *m* independent attention mechanisms are integrated to achieve specific embedding:

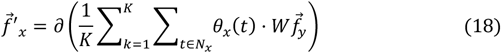

### 2.5 Method overview

#### GAPDA

In this study, we propose a novel method called GAPDA to predict biologically significant, yet unmapped associations between piRNA and disease on a large scale. GAPDA is generally composed of five components, the process shown in Figure 5. First, we construct piRNA and disease feature descriptors based on sequence information, disease semantic information, and Gaussian interaction profile kernel similarity information. Therefore, the final feature 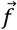 is defined as follows:

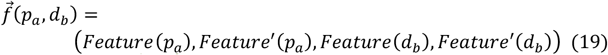

**Fig. 5.**
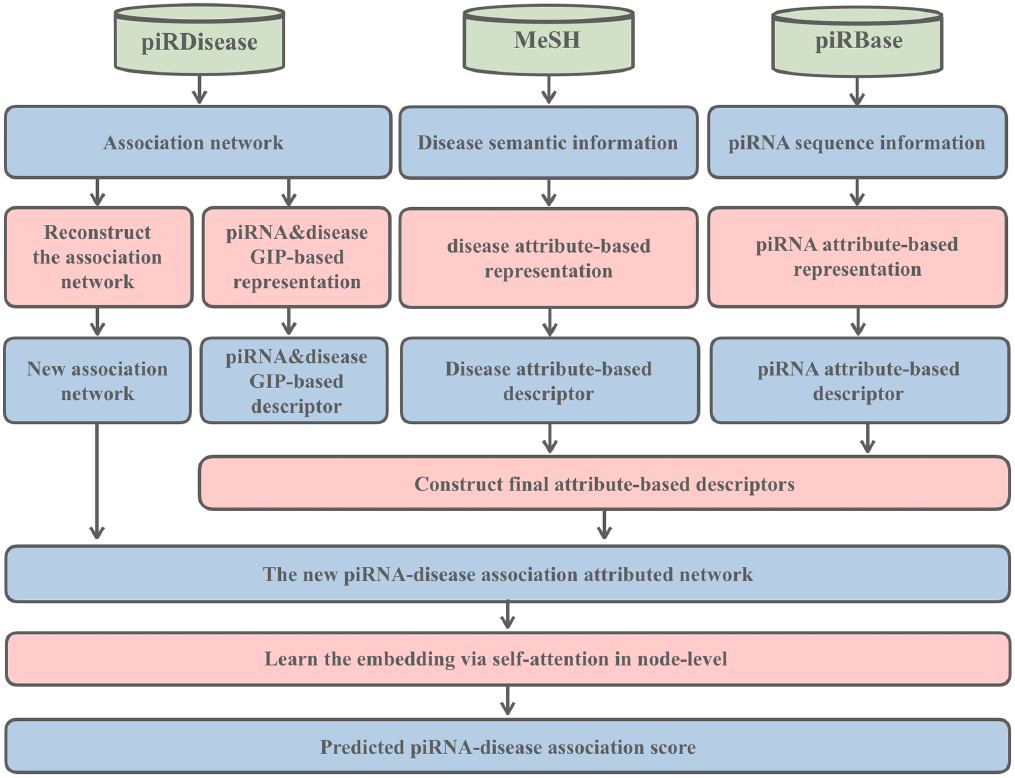
The flowchart of GAPDA for predicting piRNA-disease association.

Second, based on the existing associated network, an abstract network topology is constructed to expand the information contained in the network. Third, the reconstructed abstract network topology is combined with the final descriptor 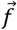 to obtain a new piRNA-disease association attribute network. Fourth, the network embedding in node-level is learned via the attention-based graph neural network. Finally, the degree of association between piRNA and disease pairs is scored. In particular, the predicted scores of piRNA and disease pairs are directly proportional to the probability of association.

## 3 EXPERIMENTAL RESULTS

### 3.1 The performance of GAPDA on the benchmark dataset

In this part, we choose 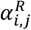 as an element for abstract network topology. In order to evaluate the performance of the proposed method, it is applied to the benchmark database GPRD.

Figure 6 depicts the ROC curve generated on the baseline data and the average AUC of five-fold cross-validation is 0.9038. In detail, the AUCs of GAPDA are 0.9115, 0.8943, 0.9109, 0.9167, 0.8859. In addition, Table 2 lists the results of the detailed evaluation criteria, with the average accuracy (Acc.) of 0.8569, the precsion (Pre.) is 0.8550, the Recall (Rec.) is 0.8638 and the F1-score is 0.8577. Their standard deviations are 0.92%, 3.56%, 4.16%, 0.92%, respectively. From the results, the lowest accuracy in the five experiments reached 0.8395, and the highest accuracy reached 0.8642. Meanwhile, this experiment relies on the network structure to make predictions, and the prediction results obtained by different attribute networks have error. Overall, our approach yielded convincing results, suggesting that GAPDA can provide powerful candidates for piRNA as a biomarker and has the potential to drive disease diagnosis and to identify disease mechanisms.

**Fig. 6.**
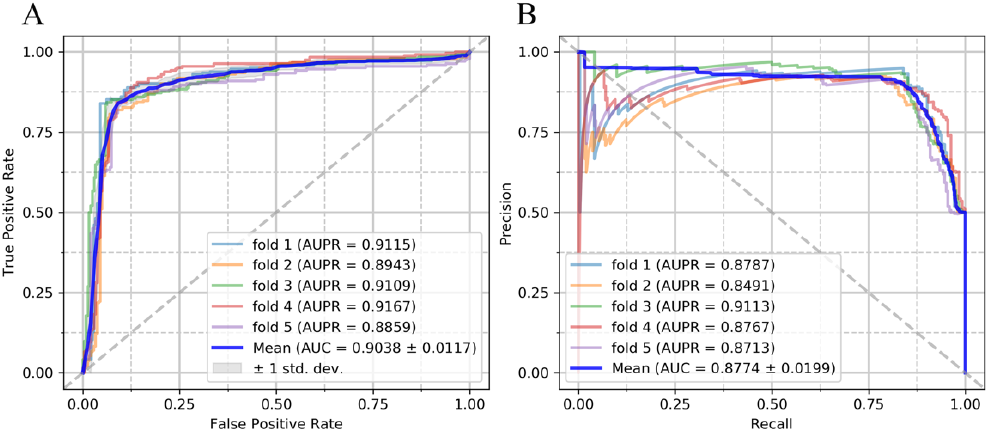
(A) ROC curves performed by GAPDA on GPRD dataset. (B) PR curves performed by GAPDA on GPRD dataset.

**Table 2.** Five-fold cross-validation results performed by GAPDA on GPRD dataset.

| Testing set | Accuracy | Precision | Recall | F1-score |
| --- | --- | --- | --- | --- |
| 1 | 0.8642 | 0.8391 | 0.9012 | 0.8690 |
| 2 | 0.8395 | 0.8022 | 0.9012 | 0.8488 |
| 3 | 0.8636 | 0.8729 | 0.8512 | 0.8619 |
| 4 | 0.8554 | 0.9095 | 0.7893 | 0.8451 |
| 5 | 0.8616 | 0.8514 | 0.8760 | 0.8635 |
| Average | 0.8569 | 0.8550 | 0.8638 | 0.8577 |
| | $\pm 0.0092$ | $\pm 0.0356$ | $\pm 0.0416$ | $\pm 0.0091$ |

### 3.2 Comparison with Attribute-based and Collaborative Filtering methods

In the association prediction model of ncRNA and disease, attribute-based (Att-based) and collaborative filtering-based (CF-based) methods are common. In order to better evaluate the performance of the proposed method, we compare it with these two methods. The results are shown in Table 3. The evaluation indicators of GAPDA are higher than the other two traditional methods, especially the accuracy. Therefore, the attention-based approach has better performance than traditional attribute-based and collaborative filtering-based approaches. In addition, other evaluation parameters are higher than the average performance. There are many reasons for the superior performance of GAPDA. First, the two traditional methods only consider attribute information or network information, and do not combine the two sources of heterogeneous knowledge. However, the proposed method combines four kinds of information into an attribute network, which can well quantify the characteristics of the association. Second, the introduction of attention mechanisms allows the hidden representation of nodes to be computed through neighbor behavior. This operation can effectively improve the performance of the model. Third, the new abstract network topology we built also helps improve performance. In the real world, networks are often heterogeneous. This method abstracts existing networks into adjacency matrix with uniform size, which is conducive to the fusion between heterogeneous networks. In addition, the results are represented as a histogram for a more intuitive comparison (Figure 7).

**Table 3.** Comparison of different types of prediction method on GPRD dataset.

| Method | AUC | AUPR | Accuracy | Precision | Recall | F1-score |
| --- | --- | --- | --- | --- | --- | --- |
| Att-based | 0.8725 | 0.8465 | 0.8200 | 0.8247 | 0.8143 | 0.8189 |
| CF-based | 0.9032 | 0.8822 | 0.8280 | 0.8329 | 0.8260 | 0.8272 |
| GAPDA | 0.9038 | 0.8944 | 0.8569 | 0.8550 | 0.8638 | 0.8577 |

**Fig. 7.**
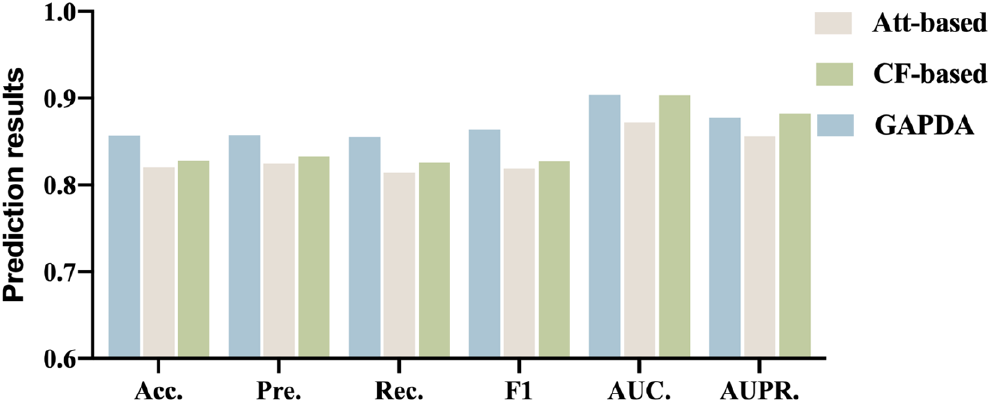
Comparison of results of Att-based, CF-based method and GAPDA on GPRD dataset.

### 3.3 Comparison of different strategies to generate abstract network topologies

In Section 2.2, an abstract network topology method to reconstruct the associated network is proposed and we design three strategies to generate an abstract network topology. In Section 3.1, the results of 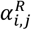 have been described. So, in this section, we evaluate the other two strategies to evaluate the performance of the abstract network topology approach. As shown in Table 4, Table 5, and Figure 8, i) based on any abstract network topology, the performance of the proposed method is higher than the average of the traditional methods. This shows that the attribute network constructed with an abstract network topology can combine multiple knowledge sources to restore the true state of the network. This can improve model performance. ii) most evaluation criteria of *A^D^* and *A^RD^* strategies are inferior to *A^R^*, of which *A^D^* strategy is the most obvious. The reason is that the elements with value=1 in the adjacency matrix *A^D^* are too dense, which makes its abstract network topology specificity insufficient, and *A^RD^* is similar. The above two information shows that different abstract network topologies affect the performance of the model to varying degrees, so giving them different weights can improve the effectiveness.

**Table 4.**
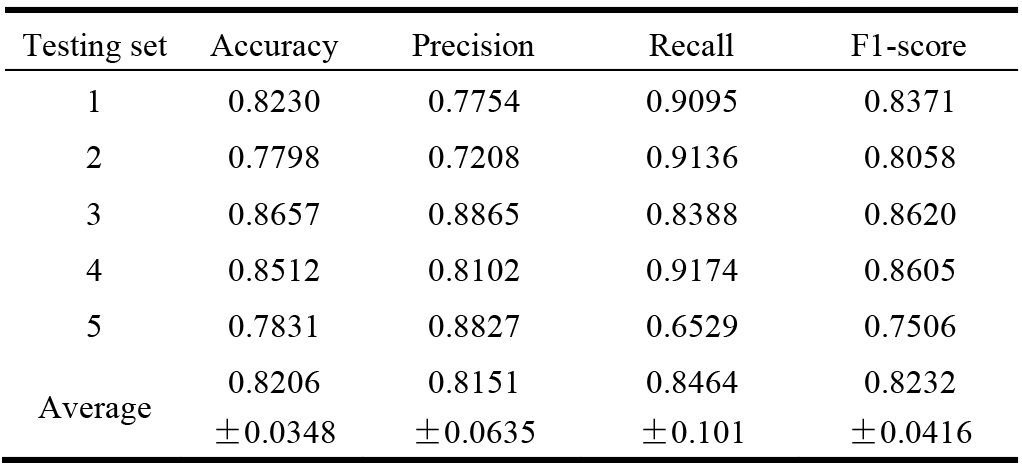
Five-fold cross-validation results performed by GAPDA (***A^D^***) on GPRD dataset.

| Testing set | Accuracy | Precision | Recall | F1-score |
| --- | --- | --- | --- | --- |
| 1 | 0.8230 | 0.7754 | 0.9095 | 0.8371 |
| 2 | 0.7798 | 0.7208 | 0.9136 | 0.8058 |
| 3 | 0.8657 | 0.8865 | 0.8388 | 0.8620 |
| 4 | 0.8512 | 0.8102 | 0.9174 | 0.8605 |
| 5 | 0.7831 | 0.8827 | 0.6529 | 0.7506 |
| Average | 0.8206<br>$\pm 0.0348$ | 0.8151<br>$\pm 0.0635$ | 0.8464<br>$\pm 0.101$ | 0.8232<br>$\pm 0.0416$ |

**Table 5.** Five-fold cross-validation results performed by GAPDA (***A^RD^***) on GPRD dataset.

| Testing set | Accuracy | Precision | Recall | F1-score |
| --- | --- | --- | --- | --- |
| 1 | 0.8807 | 0.9004 | 0.8560 | 0.8776 |
| 2 | 0.8395 | 0.8022 | 0.9012 | 0.8488 |
| 3 | 0.8368 | 0.8147 | 0.8719 | 0.8423 |
| 4 | 0.8533 | 0.8132 | 0.9174 | 0.8621 |
| 5 | 0.8182 | 0.7831 | 0.8802 | 0.8288 |
| Average | 0.8457<br>$\pm 0.0208$ | 0.8227<br>$\pm 0.0404$ | 0.8853<br>$\pm 0.0217$ | 0.8519<br>$\pm 0.0167$ |

**Fig. 8.**
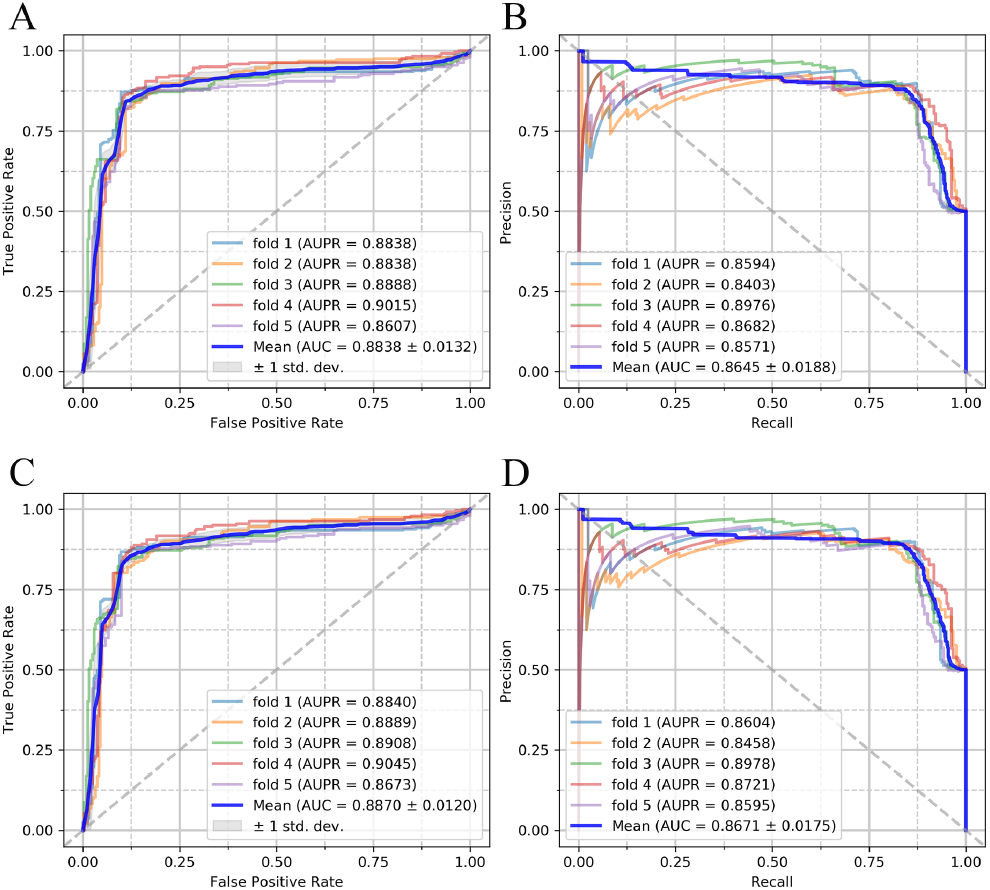
A) ROC curves performed by GAPDA (***^D^***) on GPRD dataset. B) PR curves performed by GAPDA (***A^D^***) on GPRD dataset. C) ROC curves performed by GAPDA (***A^RD^***)on GPRD dataset. D) PR curves performed by GAPDA (***A^RD^***) on GPRD dataset.

## 4 CONCLUSION

Since the network of interactions between molecules in the real world is enormously intricate and noisy, how to efficient graph mining becomes a hot spot. In this study, we propose a piRNA-disease association prediction framework based on the graph attention network to capture graph features and calculate the hidden representations of associations in the network based on neighbor nodes. In particular, we introduced attention-based graph neural networks into the field of bio-association prediction for the first time, and proposed an abstract network topology suitable for small samples. Supported by these two novel methods, GAPDA showed encouraging results in predicting piRNA-disease associations. In detail, in the five-fold cross-validation, GAPDA got an AUC of 0.9038, AUPR of 0.8774, and accuracy of 0.8569. In addition, we compared two traditional methods and different strategies to generate abstract network topologies. Experiments showed that GAPDA can be an excellent complement to future biomedical research and has determined the prospect of the graph neural grid on such problems. We hope that the proposed method can provide a powerful candidate for piRNA biomarkers and can be extended to other graph-based tasks.

## ACKNOWLEDGEMENTS

We thank anonymous reviewers for very valuable suggestions.

